# Theta-burst direct electrical stimulation remodels human brain networks

**DOI:** 10.1101/2024.02.15.580568

**Authors:** Yuhao Huang, Rina Zelmann, Peter Hadar, Jaquelin Dezha-Peralta, R. Mark Richardson, Ziv M. Williams, Sydney S. Cash, Corey J. Keller, Angelique C. Paulk

## Abstract

Patterned brain stimulation is a powerful therapeutic approach for treating a wide range of brain disorders. In particular, theta-burst stimulation (TBS), characterized by rhythmic bursts of 3-8 Hz mirroring endogenous brain rhythms, is delivered by transcranial magnetic stimulation to improve cognitive functions and relieve symptoms of depression. However, the mechanism by which TBS alters underlying neural activity remains poorly understood. In 10 pre-surgical epilepsy participants undergoing intracranial monitoring, we investigated the neural effects of TBS. Employing intracranial EEG (iEEG) during direct electrical stimulation across 29 stimulation cortical locations, we observed that individual bursts of electrical TBS consistently evoked strong neural responses spanning broad cortical regions. These responses exhibited dynamic changes over the course of stimulation presentations including either increasing or decreasing voltage responses, suggestive of short-term plasticity in the amplitude of the local field potential voltage response. Notably, stronger stimulation augmented the mean amplitude and distribution of TBS responses , leading to greater proportion of recording sites demonstrating short-term plasticity. TBS responses were stimulation site-specific and propagated according to the underlying functional brain architecture, as stronger responses were observed in regions with strong baseline effective (cortico-cortical evoked potentials) and functional (low frequency phase locking) connectivity. Further, our findings enabled the predictions of locations where both TBS responses and change in these responses (e.g. short-term plasticity) were observed. Future work may focus on using pre-treatment connectivity alongside other biophysical factors to personalize stimulation parameters, thereby optimizing induction of neuroplasticity within disease-relevant brain networks.

## Introduction

Transcranial magnetic stimulation (TMS) represents a leading candidate as a circuit-based intervention to treat dysfunctional brain circuits in psychiatry and neurology even including stroke ^1–4^. Indeed, about half of patients with depression who have not responded to medication demonstrate a clinical response to TMS (reduction in symptoms by ≥50%)^2,3^. Further, patterned TMS such as *theta burst stimulation* (or *TBS*), has shown promise as a clinical tool. TBS involves high frequency (50-200 Hz) bursts spaced at theta rhythm (3-8 Hz) ^5–14^. Recently, daily noninvasive TMS-delivered TBS to the left dorsolateral prefrontal cortex (DLPFC) ^5–7,15^ was FDA-cleared for treatment of medication-resistant major depressive disorder ^16^. Indeed, because TBS is much shorter in time than the previously-standard 10 Hz protocol (3 minutes versus 37.5 minutes, respectively) ^5^, TBS is now becoming standard in the field. Further, instead of once daily TBS treatments, ‘accelerated’ TMS-delivered TBS approaches have been developed -- and for some forms, now FDA-cleared to treat depression -- whereby up to ten TMS-delivered TBS treatments are delivered daily ^17–19^.

Despite these exciting innovations and the potential for personalized target selection as well as increasing access to care ^20–22^, TMS clinical trials show widely variable results. Further, clinical response one month after treatment remains at 50% ^5,17,23,24^. Factors contributing to treatment heterogeneity include a large treatment parameter search space and incomplete understanding of neural effects. An improved understanding of the neural mechanisms underlying how TBS directly alters brain activity can reveal pathways to improve treatment efficacy.

The clinical efficacy of TMS likes hinges on factors such as pattern, timing and location, although empirical neural evidence in humans is scarce. Recent studies indicate that repetitive TMS not only modulates neural firing ^9,25,26^, but also induces enduring changes in neural activity patterns over several minutes ^27^, suggesting a mechanism of neuroplasticity underlying its clinical effects ^27–30^. Indeed, the rationale behind employing TBS-patterned stimulation in clinical treatment stems from its demonstrated efficacy in inducing long-term potentiation in slice physiology and rodent models ^31–37^. Furthermore, the cumulative effect of TMS treatment across multiple sessions has been substantiated by preclinical data in the motor cortex, showing enhanced and prolonged changes in excitability with multiple sessions compared to single sessions ^11,38–40^ . Moreover, stimulation location is likely important for effective treatment, given motor TMS for stroke ^4^ or the left DLPFC TMS for depression ^16^ are needed in achieving specific therapeutic outcomes. In summary, inducing neural changes that relate to clinical outcome may depend considerably on stimulation location, intensity, and timing. Yet, elucidating these relationships on the neural level continues to prove challenging in the non-invasive space such as with fMRI, EEG. We propose uncovering the neural mechanisms underlying TBS-patterned stimulation and its effects on neural activity by using focal patterned direct electrical stimulation coupled with high spatiotemporal resolution intracranial brain recordings.

Direct electrical stimulation via intracranial leads has been a mainstay in modulating neural activity to uncover brain function and treat neurological and psychiatric disorders. Research has focused on developing personalized neuromodulatory therapies via deep brain stimulation (DBS) have led to significant progress in optimizing stimulation parameters such as current, frequency, and patterns in a tailored fashion ^41–53^. This has translated to clinical success, whereby DBS is used for treatment of neuropsychiatric conditions, notably medication-resistant major depressive disorder and OCD ^17,54,55^. As such, intracranial stimulation coupled with intracranial EEG (iEEG) is emerging as a powerful method to study the mechanisms of TBS. This approach provides anatomically precise information about neuronal populations at a millimeter scale and temporally precise information on neural dynamics at a millisecond scale. Direct electrical TBS-patterned stimulation paired with iEEG measurements have found lasting entrainment of frequency-specific oscillations (in the theta band, 4-8 Hz) after TBS ^12^, TBS-specific induced buildup of beta band coherence in the sensorimotor cortex ^56^, and improvement in memory within a learning and memory task via TBS microstimulation of medial temporal lobe structures ^8^. Crucially, the location of the TBS-specific responses were highly correlated with brain functional connectivity to stimulation ^12^. Hence prior studies have demonstrated that intracranial TBS elicits neural changes persisting beyond mere seconds, shaped by the underlying functional connectivity in the brain ^12,56^. However, these studies predominantly focused on oscillatory changes, overlooking potential effects on voltage responses both within and across the TBS train (in the interval between bursts) or across stimulation trains (between rounds of TBS bursts). While non-human studies, such as those examining long term potentiation (LTP) ^57^ have historically examined the temporal evolution of voltage responses, equivalent investigations in humans have been sparse, particularly in brain regions pertinent to psychiatric disorders. Moreover, there remains a paucity of research addressing how neural changes unfold in response to repeated patters of direct electrical TBS in intracranial human brain recordings ^58–60^. Closing these gaps in understanding promises to deepen our grasp of the neural mechanisms underpinning TBS and its implications for neuropsychiatric disorders.

In an effort to elucidate the neurophysiological mechanisms underlying TBS, we evaluated the neural effects of electrically-delivered focal TBS as a function of location, time, and amplitude using iEEG voltage recordings in participants with medically-intractable epilepsy. We chose stimulation locations that in prior studies demonstrated stimulation-induced memory or neuropsychiatric improvements in individuals, such as the anterior cingulate cortex (ACC), or temporal lobe cortex, ventrolateral prefrontal cortex (VLPFC), and dorsolateral prefrontal cortex (DLPFC) ^1,61–67^. Given TBS’s purported ability to induce persistent neural changes^39,56,68^, we hypothesized that TBS triggers discernible signatures of changing voltage responses both within and across repeated trains of intracranial TBS. We further posited that these changes are influenced by stimulation intensity, location specificity, and the baseline functional, structural, and effective connectivity of the targeted brain network ^13,43,50,69–76^. Our findings revealed robust voltage responses to single TBS bursts with these acute responses displaying dynamic changes in amplitude both within and across trains of stimulation which include both increasing and decreasing responses over stimulation presentations, suggesting a form of short-term plasticity. Notably, these response dynamics were dependent on stimulation intensity and targeted brain network. Furthermore, we demonstrated the predictive ability of baseline connectivity measures in forecasting TBS response patterns. Collectively, these results underscore the capacity of TBS-patterned electrical stimulation to induce region-specific short-term plasticity in the human brain.

## Materials and Methods

### Human Participants and Recordings

We recorded intracranial neural activity from 10 participants with intractable epilepsy undergoing evaluation through invasive monitoring. In all cases, participants underwent stereo-electroencephalography, with implantation of multi-contact depth electrodes to locate epileptogenic tissue in relation to essential cortex (**Supplemental Table 1**). Depth electrodes (PMT, Chanhassen, MN, USA) with diameter 0.8 and 4-16 platinum/iridium-contacts (electrodes) 1-2.4 mm long with inter-contact spacing ranging from 4-10 mm (median 5 mm) were placed stereotactically, based on clinical indications for seizure localization determined by a multidisciplinary clinical team independent of this research. Following implant, the preoperative T1-weighted MRI was aligned with a postoperative CT using volumetric image coregistration procedures and FreeSurfer scripts ^77–82^ (http://surfer.nmr.mgh.harvard.edu). Electrode coordinates were manually determined from the CT in the patients’ native space ^80,81^ and mapped using a surface based electrode labeling algorithm (ELA; ^80–82^) and a volume based electrode volume labeling approach ^81^ that registered each contact to the DKT atlas ^83^. To map the electrode locations to common brain locations in MNI (Montreal Neurological Institute) space, we used MATLAB and Fieldtrip tools (http://www.ru.nl/neuroimaging/fieldtrip) ^84^. Surface representation of quantified measures on a common pial surface was performed using the ECoG/fMRI visualization and data plotting toolbox for Matlab toolbox (https://edden-gerber.github.io/vis_toolbox/ ; plot_data_on_mesh.m; patch size = 30; overlap method = mean).

In all cases but one, participants had received their normal antiepileptic medications prior to stimulation to minimize the risk of seizure. Recordings used a Blackrock system with FrontEnd amplifiers with a sampling rate of 2 kHz (Blackrock Microsystems, Salt Lake City, UT, USA). Depth recordings were referenced to an EEG electrode placed on skin (C2 vertebra or Cz or mastoid scalp electrode) or a chest EEG surface contact.

For a subset of participants (N=4), neural activity during single pulse electrical stimulation (SPES, **Supplemental Table 1**) has been presented in previous publications with different analyses ^85,86^.

### Ethics statement

All patients voluntarily participated after fully informed consent as monitored by the Massachusetts General Brigham Institutional Review Board covering Massachusetts General Hospital (MGH). Participants were informed that involvement and engagement in the stimulation tests would not alter their clinical treatment in any way, and that they could withdraw at any time without jeopardizing their clinical care. Electrode placement and anatomical localizations were placed for seizure localization determined by a multidisciplinary clinical team purely for clinical indications and that research participation played no role in the decisions for electrode placement.

### Direct electrical stimulation

We applied direct electrical stimulation to consecutive contact pairs (bipolar pairs) across the brain. We targeted frontal and lateral temporal lobe regions in or near the gray-white matter boundary as direct electrical stimulation has been shown to engage both local circuits and distant brain networks ^85^. Stimulation locations were originally chosen from sites in or near the anterior cingulate cortex (ACC) or lateral temporal lobe cortex, or the ventrolateral prefrontal cortex (VLPFC) and dorsolateral prefrontal cortex (DLPFC) based on MRI colocalization of channels ^1,61–67^. In some cases (N=7), we performed single pulse electrical stimulation (SPES) in several sites (mean 22.1 ± 8.3 STD sites per participant) and SPES responsiveness along with location in those sites was used to inform the location of TBS delivery. Specifically, within a specific region we prioritized contact pairs that showed response to SPES outside of the local region, since we were interested in downstream activity ^85^ . We also chose electrode locations outside of areas of seizure onset as judged by the monitoring clinicians. Sites near the corpus callosum as well as sites which were too medial in the cingulate were not used to avoid potential discomfort from direct dural stimulation. Regarding the brain region classifications, we had originally only targeted the VLPFC, ACC, DLPFC, and lateral temporal lobe, but, due to contact availability and sites chosen based on a visual inspection of the coregistered channel locations on the MRI, we chose some sites that were then automatically classified to postcentral or OFC or insula. As such, we kept those sites classified as separate from the other regions.

A CereStim stimulator (Blackrock Microsystems, Salt Lake City, UT) was used to deliver TBS or SPES stimulation. Current injection and return paths used neighboring electrodes in a bipolar configuration ^41,53^. Stimulation was controlled via a custom CereStim API via MATLAB or a custom C++ code (https://github.com/Center-For-Neurotechnology/CereLAB). For both TBS and SPES, the same 233 µs duration waveforms were used: 90 µs charge-balanced biphasic symmetrical pulses with an interphase interval of 53 µsec at 7 mA between 10 and 25 trials (for SPES) ^69,74,85–90^ and TBS at 1 mA and 2 mA. The interval of 53 µs within the biphasic waveform was required as a hardware-limited minimum interval between square pulses with the CereStim stimulator. While SPES involved only one set of the charge-balanced biphasic symmetrical pulses spaced 4-5 seconds and at different locations, TBS involved stimulating with five bursts of 200 Hz stimulation across 50 ms (which involved 10 charge-balanced biphasic symmetrical pulses). Burst were spaced 117 ms apart. We chose 200 Hz as the frequency of each burst as it has been shown to produce more consistent responses across brain regions and individuals ^42,44,45^. Five sequential bursts were used to keep the per-trial total duration of stimulation closer to 0.5 sec and to mimic previous publications^12,42^. The five bursts together were considered a single ‘train’. Then, ten trains (each with 5 bursts of 200 ms trains, with 10 trains chosen for the sake of time and to be able to stimulate multiple sites per participant) were spaced 20 seconds apart with a jitter in timing pulled from a random distribution of time with a maximum of ± 2 seconds. First, ten trains were performed at a single bipolar pair at 1 mA. Next, usually around a minute later, another ten trains at the same bipolar pair at 2 mA were performed if the monitoring epileptologist allowed it and there was no reported sensation from the participant. In total, 29 total unique sites were stimulated across participants. Five of these bipolar pairs had TBS stimulation at 1 mA only while all other sites had both 1 mA and 2 mA stimulation. All ten participants had both SPES and TBS stimulation testing at the same (s=25), or neighboring (s=4) bipolar pairs. Across all ten participants, a median of 2.5 sites were stimulated with TBS (**Supplemental Table 1**). Responses were recorded in a total of 4567 clinically implanted bipolar-referenced channels across participants. Channels, in this case, are defined as the bipolar-referenced signal from pairs of electrodes.

A trained electroencephalographer monitored ongoing recordings for epileptiform activity and asked participants if they experienced any sensations. In only one case did a participant report some sensation at which point we stopped stimulation and did not use data from stimulation at that site in our analyses. We never had to stop TBS or SPES stimulation for clinical reasons such as inducing epileptiform or seizure-like activity. Otherwise, the participants were awake and were aware that they were being stimulated but were blind to the stimulation timing and parameters.

### Electrophysiologic Analysis

Data analysis was performed using custom analysis code in MATLAB and Fieldtrip (http://www.ru.nl/neuroimaging/fieldtrip) ^84^. Channels with excessive line noise or without clear neural signal (determined by visual inspection) were removed from the analysis. The remaining electrodes were demeaned and bipolar re-referenced relative to nearest electrode neighbors to generate a signal represented on the channel level ^91–93^.

### Resting Phase Synchrony to measure Functional Connectivity

To estimate functional connectivity through oscillatory synchrony of two brain regions, we computed phase locking value (PLV) between all possible electrode pairs using FieldTrip (ft_connectivityanalysis) ^84,91,94^. PLV provides a measure of inter-regional synchrony based on phase difference between the paired signals. To calculate PLV, we divided the resting, pre-stimulation period ranging from 2 to 10 minutes into 2s epochs. We used a 4^th^ order Butterworth filter to obtain the analytical signal between 5-13 Hz, and performed Hilbert transform to obtain the instantaneous phase of the signal. The phase difference between the two signals is used to compute PLV. We chose a low-frequency range (5-13 Hz) to assess functional connectivity as it has been previously used to measure functional connectivity in the setting of studying TBS ^13^. As a separate measure of resting connectivity, we also computed the Pearson’s correlation between the voltage trace of pairwise channels in the same 2s epochs. The correlation was subsequently averaged across the entirety of the resting data recording.

### CCEP Mapping to measure Stimulation-Induced Effective Connectivity

To examine effective connectivity of the stimulated network, we used responses to SPES to perform corticocortical evoked potential (CCEP) mapping ^43,44,50,69,72,74,85–90,95^. Since we were comparing the SPES-induced CCEP responses with the TBS responses, we applied SPES in the same or nearby contact pairs as the TBS stimulation. Only 4 sites had SPES applied in a neighboring pair of sites compared to TBS, whereas the remaining 25 sites had SPES and TBS applied to the same region. A median of 20 SPES trials (range: 4 to 40) were applied. As stated above, if a site induced a sensation, we stopped stimulation at that site (which occurred in 2 SPES sites). If a site induced sensation with SPES, we did not use that site for TBS. CCEPs from each channel were first epoched from -1000 to 1000 ms. The epoch was subsequently standardized using Z-scores against the pre-CCEP baseline period (-150 to 1000 ms; **Figure 1D Panel 2**). We used a basis profile curve parameterization approach to quantify the mean CCEP amplitude as well as the mean CCEP duration ^96^ . Whereas conventional CCEP quantification can be dependent on the CCEP waveform and selection of the quantification time, this approach utilizes a machine learning framework that allows for general quantification of CCEP responses, irrespective of the shape of the response.

**Figure 1:**
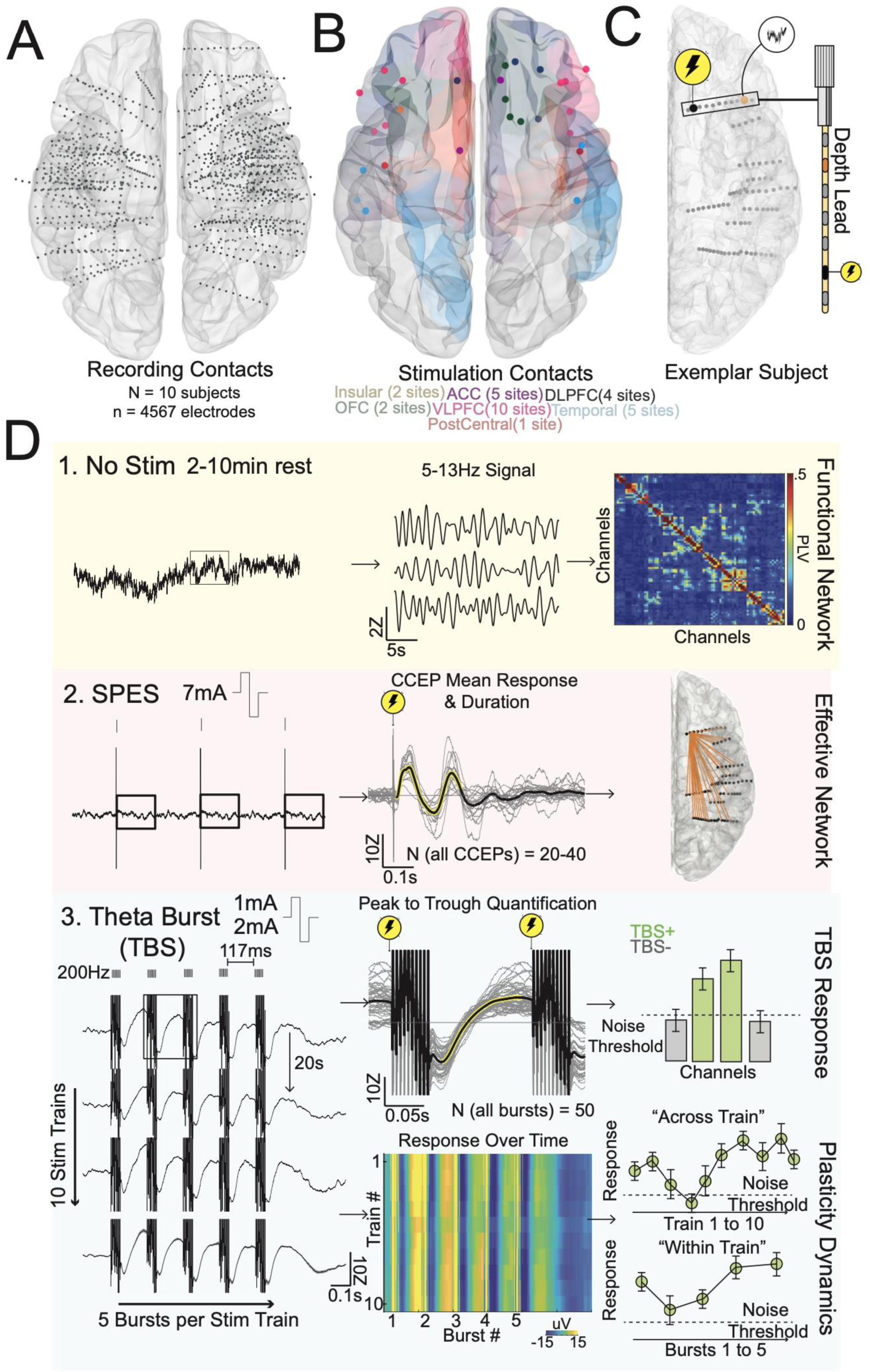
Experimental paradigm and analysis procedure. **(A)** 10 participants were enrolled with a combined total of 4567 bipolar-referenced channels. **(B)** Stimulation sites were selected across cortical regions including the anterior cingulate (ACC, 5 sites), the postcentral gyrus (1 site), the dorsolateral prefrontal cortex (DLPFC, 4 sites), the orbitofrontal cortex (OFC, 2 sites), the ventrolateral prefrontal cortex (10 sites), the insular cortex (2 sites), and the lateral temporal cortex (5 sites). **(C)** Trials of single pulse electrical stimulation (SPES) and theta-burst stimulation (TBS) were delivered at specific sites during each experimental session, while continuous iEEG was obtained at all other channels. **(D)** Schematic representation of analyses performed. Resting iEEG data (3-10 minutes) prior to stimulation was used to construct a functional network using low-frequency amplitude and phase coupling. From SPES, cortical-cortical evoked potential (CCEP) mean and duration were quantified through parameterization to estimate effective connectivity. Lastly, theta burst stimulation (TBS) consisted of ∼4 minutes of 10 stimulation trains, with each train consisting of five theta-frequency bursts, separated by 20s. Stimulation amplitude was applied first at 1 mA and then at 2 mA sequentially. TBS mean response was defined as the peak-to-trough response post-burst across all bursts (N = 50 bursts). Channels with TBS mean post-burst response above noise threshold are considered significant (TBS+). In TBS+ channels, successive post-burst responses are analyzed with reference to the train number or the burst number. Repeated-measures ANOVA was used to determine if there was a significant burst number effect (*within train* plasticity) or a significant train number effect (*across train* plasticity). Figures in **D** are for visualization and schematic purposes only.

### Theta-Burst Stimulation Response and Dynamics Quantification

To examine cortical responses to TBS, we first aligned the iEEG signal to the start of each theta-burst and epoched the data -1s to 1s to the offset of the burst, producing 50 observations (5 theta-bursts per train * 10 trains; **Figure 1D Panel 3**). Each observation or epoch was subsequently standardized using Z-scores against a pre-train baseline (-2s to -1.5 prior to the start of each train). A pre-train baseline period was chosen because the pre-burst time period, which is part of the previous burst’s post-burst response, has not returned to ‘resting’ voltage levels. The post-burst evoked response was quantified by taking the peak-to-trough amplitude from 0.01s to 0.1s post-burst. The 10ms delay was to avoid contamination from the stimulation artifact, which had a rapid drop off by 2ms when we aligned each epoch to the offset of the burst stimulation. To compare to the post-burst response, a ‘baseline’ response was also quantified, by measuring the peak-to-trough amplitude from -0.1s to -0.1s (as a measure of maximum variance) prior to the start of each train. A two-sample t-test was used to compare the post-burst evoked response against the pre-train ‘baseline’ response. The post-burst evoked response was considered significant at alpha of 0.05, after FDR correction for multiple channels comparison. A channel with a significant post-burst evoked response is abbreviated as TBS+. To evaluate the temporal dynamics of the post-burst evoked response as a function of burst order, we used a repeated-measures ANOVA (**Figure 1D; Panel 3**).

To quantify the dynamic nature of the TBS response, we define *short-term LFP plasticity* (referred to as plasticity in this manuscript) as a significant change in post-TBS response either across TBS bursts within train or across trains. Only channels that demonstrated significant post-burst evoked responses as defined above were considered for analysis of *change in the TBS response*, indicating plasticity. This thresholding procedure (only using channels which exhibited strong TBS responses at the single burst level), was to ensure the response being studied over time reflects a change in the underlying amplitude of the evoked response, and not due to a drift in a weak post-burst signal. In the ‘burst’ or ‘within-train’ dimension (five consecutive bursts delivered at theta frequency; see above), we fitted a repeated measures ANOVA model whereby bursts 1 to 5 are the repeated measures, and stimulation train 1 to 10 is the predictor variable (Burst Responses ∼ Stimulation Trains). A channel was considered to have demonstrated ‘**within-train plasticity**’ if the coefficient for the repeated measure was significant at an alpha of 0.05 after FDR correction for multiple channels comparison. Similarly, in the stimulation train dimension, we fitted a repeated measures model whereby stimulation trains 1 to 10 are the repeated measures, and burst order 1 to 5 is the predictor variable (Stimulation Train Responses ∼ Burst Order). A channel was considered to have demonstrated ‘**across-train plasticity**’ if the coefficient for the repeated measure was significant at an alpha of 0.05 after FDR correction for multiple channels comparison.

### Binary Classification Analysis

To determine if baseline structural and functional features can be used to predict TBS responses on the per-channel basis, we used multivariate logistic regression. Three sets of data were used. 1) To predict spatial distribution of significant post-burst responses, all channels across participants and sessions were pooled, and categorized by presence or absence of a significant post-burst response. 2) To predict presence of TBS-induced plasticity, only channels with significant post-burst responses were selected, and are categorized by presence or absence of plasticity (with-train or across-train plasticity). 3) To predict the type of plasticity, only channels demonstrating TBS-induced plasticity were used, and were categorized as within-train plasticity, across-train plasticity, or both forms of plasticity. Channel subselection was done for plasticity prediction (dataset 2 and 3) since only channels with significant post-burst responses were considered for analysis of plasticity. To control for possible effects of pure volume conduction, we further stratified channels as being local (<30mm) or distant (>30mm) to the stimulation site. Features used in the classification analysis included both structural and functional metrics. These included proximity to white matter, Euclidean distance to the stimulation site, CCEP amplitude, CCEP duration, PLV, and voltage correlation. Using these five features, we performed logistic regression with ten-fold cross validation. Receiver operating characteristic (ROC) curves were constructed, and we quantified the area under the curve (AUC) to evaluate model performance. Mean AUC and 95% confidence interval were constructed based on variance from the ten-fold cross validation scheme.

### Statistical analysis

Binary group comparisons were done using Mann-Whitney test or two-sample t-test for independent samples and signed rank test or one-sample t-test for paired samples. The decision to use parametric tests was made when approximate normal distribution of data for the comparison groups was observed. For comparison of more than two groups, we performed ANOVA tests or Kruskal Wallis tests. K-means clustering was performed on dynamics of post-burst responses across either the burst or the train dimension. To determine the optimal number of clusters, we evaluated four clustering criteria which includes the Calinskin-Harabasz method, DaviesBouldin method, gap method, and the silhouette method (evalclusters.m; Matlab 2022b). If the evaluation criterions did not converge at an optimal cluster number, we did not perform clustering and instead evaluated the approximate direction of change across channels. For across train response patterns, the criterions did not converge. To assess approximate trends in the channels with across train plasticity, we compared post-burst responses in the first two trains (5 bursts per train; n = 10 bursts) against post-burst responses in the last two trains (n = 10 bursts) using the Mann-Whitney test. Channels demonstrating a positive trend were identified with P < 0.05 and a Mann-Whitney Z-value that was positive, whereas channels with a negative trend were identified with P < 0.05 and a Mann-Whitney Z-value that was negative. In preliminary analyses, the total channels sampled (n = 8540) is the total channels sampled reflects the total number of stimulation sessions (**Supplemental Table 1**).

### Data and Code Sharing Statement

Custom Matlab code (version R2022b) and python code in combination with open source code from the Fieldtrip toolbox (http://www.fieldtriptoolbox.org/) was used for the majority of the neural data preprocessing and analyses, with the code shared on Github (https://github.com/KellerLab-Stanford/Analysis-TBSiEEG). Stimulation was controlled via a custom CereStim API via MATLAB or a custom C++ code (https://github.com/Center-For-Neurotechnology/CereLAB). Reconstruction of electrode locations was done using the open source, free software Freesurfer (https://surfer.nmr.mgh.harvard.edu/) and MMVT (https://github.com/pelednoam/mmvt) along with MATLAB code GitHub page (https://github.com/Center-For-Neurotechnology/Reconstruction-coreg-pipeline) and detailed in the online protocol ^81,97^. Violin plots showing the distribution of the data were produced using code by Zhaoxu Liu (2023) violin plot (https://www.mathworks.com/matlabcentral/fileexchange/120283-violin-plot-and-ggtheme?s_tid=srchtitle) on the MATLAB Central File Exchange (Retrieved April 15, 2023). Activity visualization was performed using the EcoG/fMRI visualization and data plotting toolbox for MATLAB toolbox (https://edden-gerber.github.io/vis_toolbox/).

Upon publication, deidentified stimulation data will be uploaded to the Data Archive BRAIN Initiative (DABI, https://dabi.loni.usc.edu/home) in the iEEG BIDS format using a modified version of the open source code (https://github.com/bids-standard/bids-starter-kit). ^98,99^

## RESULTS

Theta-burst stimulation (TBS) induced voltage dynamics were recorded from 10 participants (median age = 27, ranging from 18 to 53 years old; five female; all but one right-handed; **Supplemental Table 1**). Participants were implanted with depth electrodes for clinical seizure monitoring, for a total of 4567 clinically implanted bipolar-referenced channels (**Fig. 1A**). Cortical sites chosen for TBS included the insula (n=2 sites), anterior cingulate cortex (ACC, n=5 sites), dorsolateral prefrontal cortex (DLPFC, n=4 sites), orbitofrontal cortex (OFC, n=2 sites), postcentral (n=1 site), ventrolateral prefrontal cortex (VLPFC, n= 10 sites), and lateral temporal lobe (n=5 sites) as identified using an automated parcellation algorithm^81^ (**Fig. 1B**). Per-region site number variability was due to the fact that site selection was from the original MRI and occurred before automatic parcellation and electrode localization. For the sake of consistency across participants, however, we defined the sites based on the parcellation for analyses. Across participants, TBS-patterned stimulation was delivered at 29 unique bipolar pairs, with a median of 2.5 bipolar pairs of stimulated electrodes per participant (ranging from 2 to 4 bipolar pairs stimulated per participant; mean ± standard deviation = 2.9 ± 1.0 sites) across all 10 participants at two current amplitude levels (1 and 2 mA) for all participants except one individual (who received only 1 mA TBS; **Supplemental Table 1**). TBS patterns included five consecutive 200 Hz bursts (50ms duration) per trial, with each trial ( or *train*) spaced by ∼20 seconds (**Fig. 1D**). For a given stimulation site, we computed *functional and effective connectivity* before TBS and the neural response to TBS across all channels (**Fig. 1D; Methods**).

### Theta-burst stimulation evokes consistent post-burst brain responses

In the neural activity following each TBS burst, we observed robust post-burst evoked responses in a subset of channels (**Fig. 2A-C**). Given that this post-burst TBS response exhibited a clear peak and trough pattern, we computed the peak to trough amplitude (**Fig. 2B**), which was significantly higher than the pre-train baseline (**Fig. 2C**; TBS+; 2-samples T-test: t(98)=31.8, P<0.001). In contrast, other channels exhibited no response (**Fig. 2D-F, see Methods**). Quantifying the post-burst TBS response across all stimulation sessions and channels we found that 14.4% (1233 / 8540) of channels demonstrated a significant post-burst TBS response across both 1 mA and 2 mA current levels (TBS+, N=10; n=29 TBS sites; **Fig. 2G**; 2-samples t-test, P <0.05 after FDR correction). Note the total channels sampled (n = 8540) is higher than the total number of channels across participants as the total channels sampled reflects the total number of stimulation sessions, since participants may undergo multiple stimulation sessions at different sites and different stimulation intensity (**Supplemental Table 1**).

**Figure 2:**
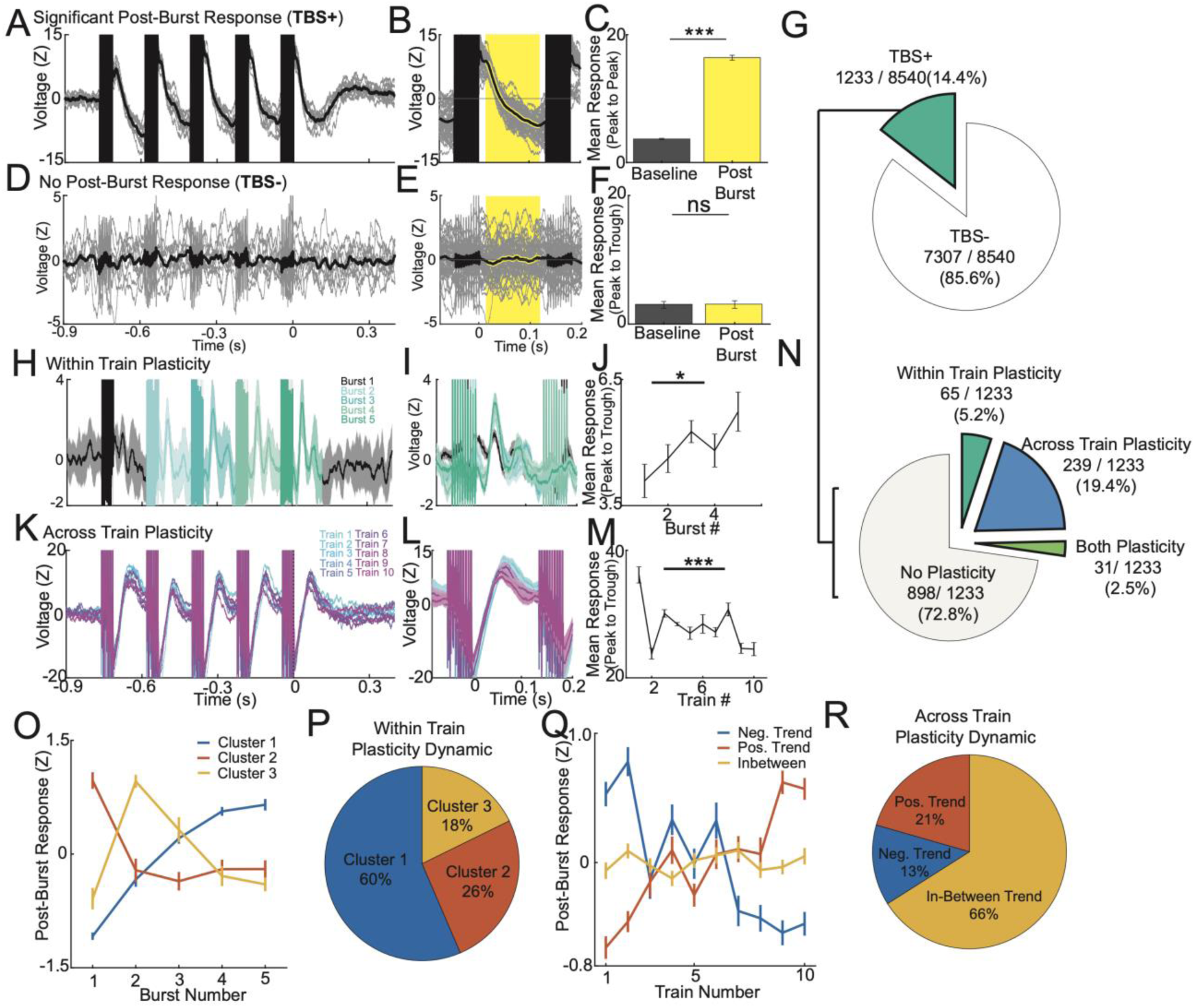
Theta-burst stimulation evokes consistent neural responses that are modulated over time. **(A,B)** Single trials and mean voltage trace in a TBS+ channel of **(A)** a train and **(B)** a single burst. Yellow shaded region denotes the post-burst quantification window. **(C)** Peak to trough quantification of the response time window in baseline and post-burst conditions. (**D,E,F**) Same as A-C but for a TBS– channel. **(G)** Among aggregate of all channels across 10 participants, 1233/8540 (14.4%) of channels were TBS+. **(H)** Mean voltage trace of a train of bursts (collapsed across trains) in a TBS+ channel exhibiting within train plasticity. Bursts 1 to 5 are highlighted in different colors. **(I)** Mean voltage trace of different bursts within the train. Note in this channel successive bursts qualitatively are larger than the first burst. **(J)** ANOVA testing of mean response across five bursts showing *within train plasticity*. **(K)** Single trial voltage traces and **(L)** mean voltage trace of the post-burst response across trains (collapsed across all bursts in a train). **(M)** Mean response across ten burst trains showing *across train plasticity*. **(N)** Among the 1233 TBS+ channels, 239 (19.4%) exhibited across train plasticity, 65 (5.2%) within train response plasticity, and 31 (2.5%) both types of plasticity. **(O)** K-means clusters of within train post-burst responses (N = 65 across 6 participants, with 41 instances during 2mA stim across multiple areas of the brain). Cluster 1 showed increasing post-burst response, cluster 2 a decrease in post-burst response, cluster 3 an initial increase and subsequently decrease. **(P)** Amongst channels with significant within train plasticity, 60% were in cluster 1, 26% in cluster 2 and 18% in cluster 3. **(Q)** K-means clustering criterion did not converge at an optimal cluster number for across train plasticity dynamics. Up-trending and down-trending post-burst responses were identified by comparing the first two trains and the last two trains. **(R)** 21% of channels (N=239) with plasticity had a positive trend, 13% had a negative trend, and 66% had a trend that did not differ in initial and final post-burst responses. Error bars reflect standard error of the mean. For all panels, *denotes P < 0.05. *** denotes P < 0.01.

### Theta-burst stimulation dynamically modulates brain responses with repeated stimulation

To quantify potential effects of TBS on short-term neuroplasticity, we asked if the post-burst response changed as a function of stimulation presentations. We define LFP plasticity as a significant change in post-TBS response either within train or across trains. In a subset of sites with significant post-burst (TBS+) responses (**Fig. 2H-M**), these responses increased with successive bursts (**Fig. 2H-J**; rANOVA: F(4,9)=2.8, P=0.037). We termed this phenomenon *within train plasticity (see Methods for details)*. In parallel, in some locations, these responses changed as a function of stimulation train (**Fig. 2K-M**; F(4,9)=15.0, P<0.001). We termed this phenomenon *across train plasticity (see Methods for details)*. Across TBS+ channels (those demonstrating a post-burst response), 19.4% (239/1233) exhibited *across train* plasticity, 5.2% (65/1233) exhibited *within train* plasticity, and 2.5% (31/1233) exhibited both types of plasticity in the combined 1 mA and 2 mA current levels (**Fig. 2N**). In the subset of channels exhibiting either form of plasticity, and to better understand the temporal dynamics, we performed k-means clustering in both the *within train* and *across train* dimensions. Multiple clustering evaluation criterions converged at three clusters as the optimal number of clusters for within train response dynamics (**Supplementary** Fig. 1A). Each cluster represented a distinct response pattern across the five bursts (**Fig. 2O**). Cluster 1 exhibited increasing responses after successive bursts, accounting for 60% of channels exhibiting within train plasticity. Cluster 2 exhibited a rapid decrease in post-burst response which persisted, accounting for 26%. Cluster 3 was characterized by an initial increase and later decrease, accounting for 18% of channels (**Fig. 2P**). K-means clustering on response dynamics across train dimension did not reach an optimal solution as different criterion diverged from each other (**Supplementary** Fig. 1B), highlighting high variability of across train response patterns. In the absence of reliable clusters, we sought to understand the proportion of channels which demonstrated either an upwards trend or a downwards trend. We compared the mean post-burst response in the first two trains against the last two trains and identified channels that showed significant difference in either direction of change (see **Methods**). We identified channels demonstrating clear uptrend or downtrend in the post-burst response across trains **(Fig. 2Q)**, which accounted for 21% and 13% of total channels with across train plasticity (**Fig. 2R**). Note that a large proportion (66%) of channels demonstrated response patterns where the initial and ending set of stimulation trains had similar post-burst responses.

### Increased stimulation current amplifies TBS-induced plasticity

We next examined how changing the stimulation current intensity influenced TBS responses to determine if there is a dose-dependent effect on either the post-burst response or on *within/across train* plasticity (**Fig. 3A**). The post-burst responses were morphologically similar, with the 2mA condition demonstrating significantly larger responses (**Fig. 3B-C**; Paired t-test: t(98)=-4.23, P<0.001). Further, while 1mA TBS led to a significant post-burst response but no plasticity, 2mA lead to both within and across train plasticity (**Fig. 3D-E**; rANOVA-Within-Train: F(4,9)=7.46, P<0.001; rANOVA-Across-Train: F(4,9)=3.12, P=0.006). At the group level, 2mA TBS led to a higher proportion of brain regions with significant post-burst responses (**Fig 3F**; 1mA median: 4.5%; 2mA median: 20%; signed rank test: Z=-4.6, P<0.001) as well as stronger post-burst responses (**Fig. 2G**; 1mA median: 3.03Z; 2mA median: 3.31Z; Z=4.29, P<0.001). With regards to plasticity, larger amplitude stimulation resulted in a higher proportion of channels that demonstrated either type of plasticity (within or across train, **Fig. 3H**; 1mA median: 1.6%; 2mA median: 3.9%; Z=-4.05, P<0.001). Specifically, this increase was driven by a higher proportion of channels exhibiting across train plasticity (**Fig.3I-K;** 1mA median: 0.7%; 2mA median: 3.0%; Z=-3.67, P<0.001). Taken together, these results indicate that higher stimulation current substantially increases cortical response to TBS, both in the proportion of post-TBS responses and dynamic modulation of the post-TBS response.

**Figure 3:**
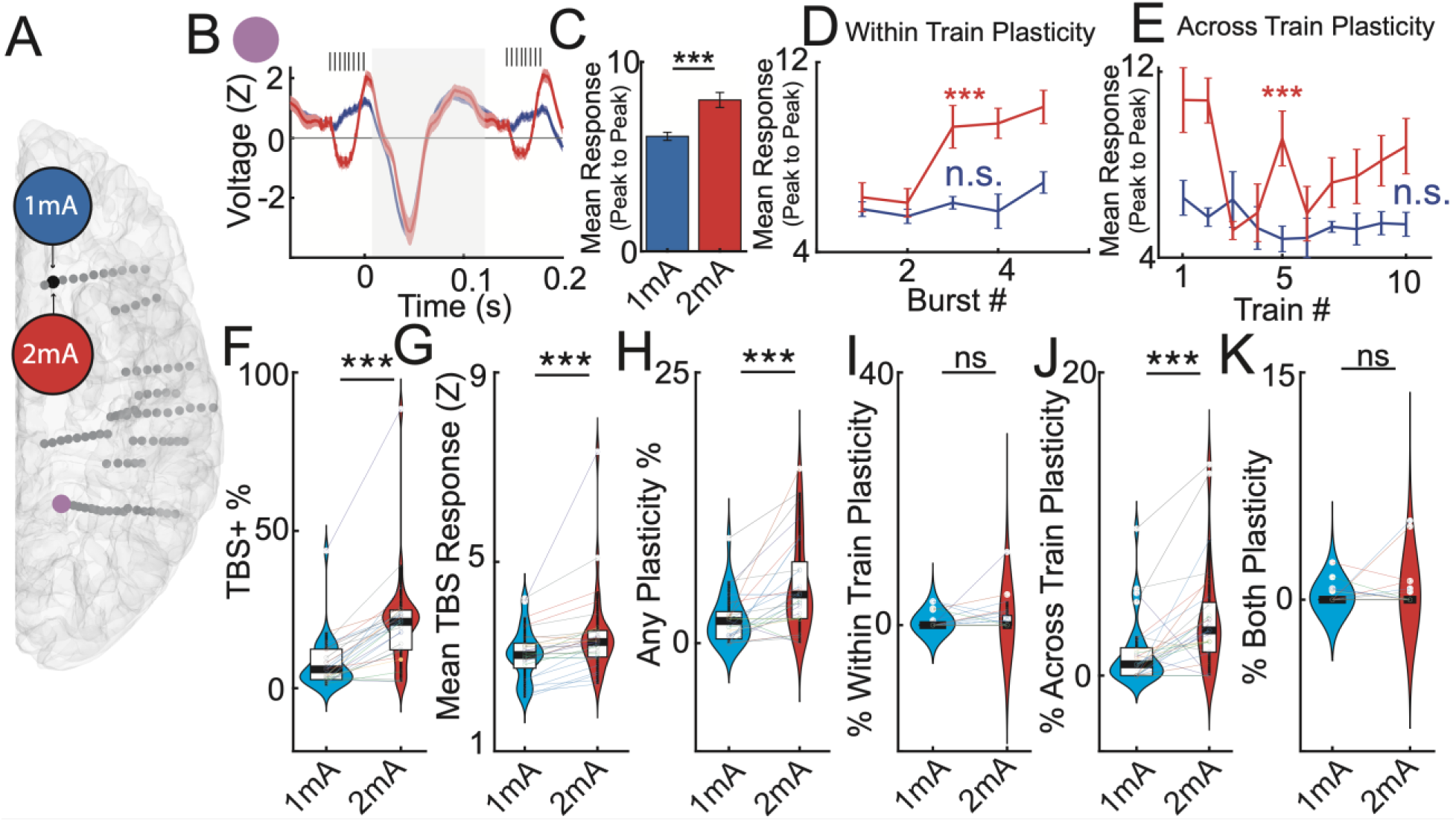
Theta-burst stimulation responses are dependent on stimulation dose. **(A)** Location of the stimulating and recording channels for **B-E**. **(B)** Mean voltage trace of the post-burst response for 1mA and 2mA stimulation. **(C)** Post-burst response was significantly different between 1mA and 2mA stimulation. **(D)** Mean response across five bursts with significant across burst plasticity noted only for the 2mA condition. **(E)** Mean response across ten stimulation trains. For each train, the five bursts are collapsed. Significant across train plasticity is noted for the 2mA condition. **(F-K) Group level effects. (F)** Proportion of TBS+ channels was significantly higher in the 2mA but not the 1mA condition. Each line represents a stimulation session, while each color represents a different participant. Higher mean post-burst response across channels **(G)** and higher proportion of TBS+ channels **(H)** with 2mA stimulation. **(I)** Proportion of channels showing within train plasticity was not different between 1mA and 2mA stimulation conditions but **(J)** proportion of channels showing across train plasticity was significantly higher. **(K)** Proportion of channels showing both types of plasticity was not different between 1mA and 2mA stimulation conditions.

### Spatial distribution of TBS responses depends on stimulation location

We next divided TBS responses based the brain region stimulated as we hypothesized that different regions would respond differentially. Across participants and combining the 1 mA and 2 mA responses, we observed that TBS produced post-burst responses in distinct regions of the brain depending on the stimulation target (**Fig. 4A**). For example, DLPFC stimulation resulted in a high proportion of significant post-burst responses in surrounding frontal regions and the cingulate. VLPFC stimulation drove responses similarly in prefrontal regions, but also included lateral and mesial inferior frontal regions. Lateral temporal lobe stimulation resulted in primarily temporal and parietal responses. Anterior cingulate stimulation resulted in significant responses in the cingulum, in the prefrontal and in parieto-occipital regions. We subsequently quantified the proportion of channels exhibiting post-burst responses (TBS+) and types of response plasticity over common anatomic divisions (**Fig. 4B**). For **DLPFC stimulation**, the top three regions exhibiting TBS responses were *ACC* (50% TBS+; 5% across train plasticity), *DLPFC* (34% TBS+; 3% across train plasticity; 1.4% within train plasticity; 1.4% both types of plasticity) and *PCC* (29% TBS+; 2% across train plasticity; 5% within train plasticity). For **VLPFC stimulation**, the top responses were *DLPFC* (28% TBS+; 4.4% across train plasticity; 3.6% both plasticity), *VLPFC* (33% TBS+; 14% across train plasticity; 1% within train plasticity; 1% both plasticity) and *OFC* (38% TBS+; 14% across train plasticity; 1.2% both plasticity). For **lateral temporal stimulation**, the top responses were *parietal cortex* (40% TBS+; 20% across train plasticity), *lateral temporal cortex* (30% TBS+; 5.6% across train plasticity; 2.6% within train plasticity; 1.2% both plasticity) and the *PCC* (23% TBS+). And for **ACC stimulation**, top responses were *DLPFC* (48% TBS+; 13% across train plasticity; 6.5% within train plasticity; 4.3% both plasticity), *PCC* (100% TBS+; 6.3% across train plasticity; 44% within train plasticity) and *occipital cortex* (52% TBS+; 6.3% across train plasticity; 4.2% within train plasticity).

**Figure 4:**
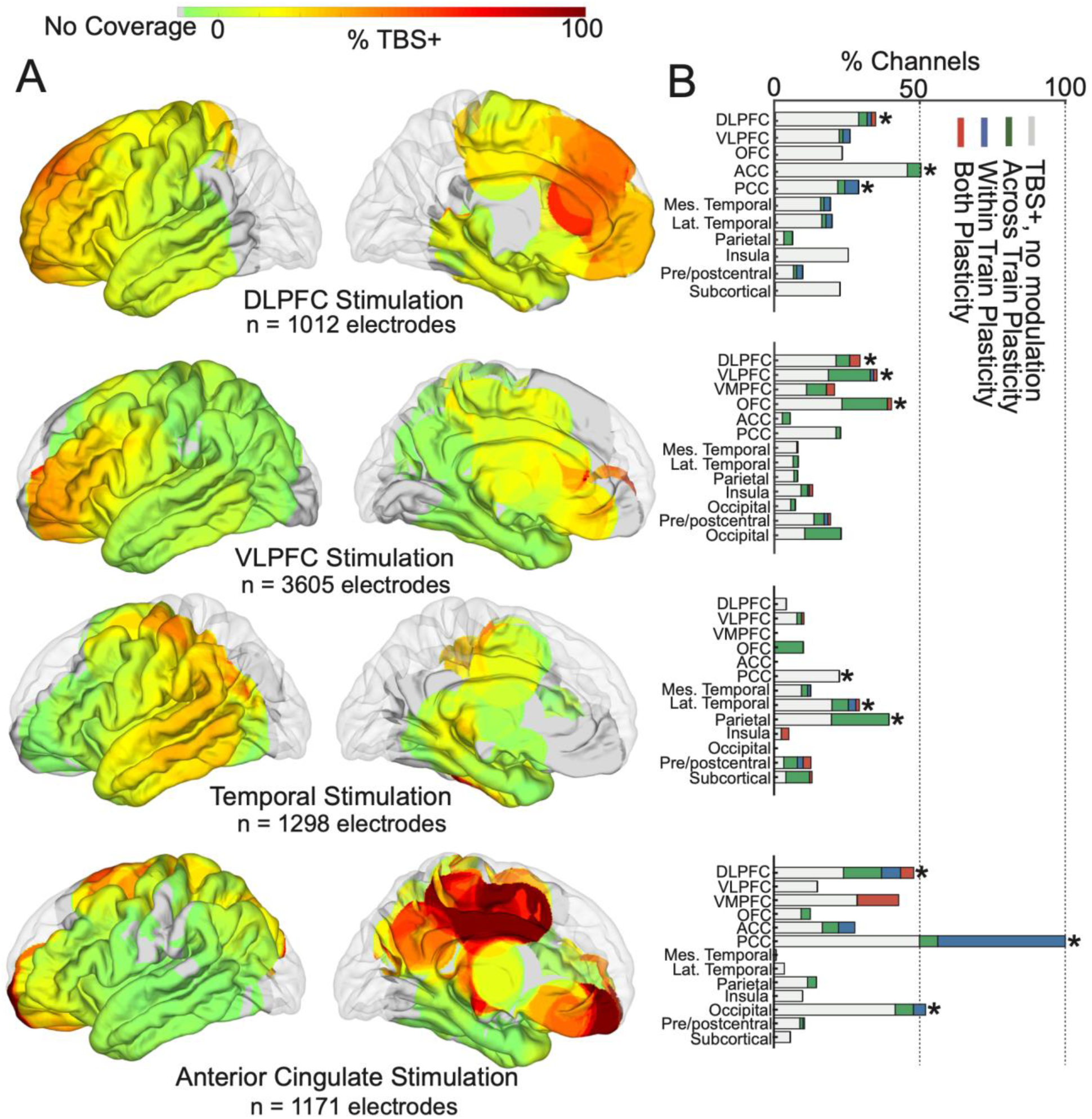
Spatial specificity of theta-burst stimulation response and plasticity by stimulation site. **(A)** Surface heatmap and bar chart **(B)** showing percentage of TBS+ local channels for a particular stimulation site within an anatomical region (gray), within train response plasticity (blue), across train plasticity (green) and both types of plasticity (red). Top three sites for each stimulation location are noted with an asterisk. Note reciprocal responses in the DLPFC and ACC when stimulated. ACC stimulation elicits widespread TBS responses. DLPFC: dorsolateral prefrontal cortex; VLPFC: ventrolateral prefrontal cortex; VMPFC: ventromedial prefrontal cortex; OFC: orbitofrontal cortex; ACC: anterior cingulate cortex; PCC: posterior cingulate cortex.

### Structural, effective, and functional connectivity constrain and predict theta-burst responses

Finally, we hypothesized that the underlying brain connectivity shapes the location and spatial extent of post-burst responses and plasticity during TBS ^13,69^. As a proxy for structural connectivity, we measured the *ratio of proximity to gray and white matter* as well as the *distance to stimulation site* for each recording channel ^13,69–71,85^. To measure other types of connectivity, phase locking value (PLV) and voltage correlations during resting state before TBS were calculated to measure functional connectivity while cortico-cortical evoked potential (CCEP) amplitude and duration were measured to evaluate stimulation-induced effective connectivity ^12,72^ (**Fig. 5A; Supplementary** Fig. 3**, see Methods for details**). Across participants and stimulation sites, we compared these structural and functional measures in regions with and without significant post-burst responses (TBS+ vs TBS-). We found that TBS+ regions (regions with significant TBS responses following stimulation) tended to reside more in white matter (signed-rank test: Z = 5.6; P<0.001), have higher functional (low-frequency PLV; Z = -6.2; P<0.001) and effective connectivity (CCEP amplitude; Z = -6.36; P < .001), and are located closer to the stimulation site (Z = 6.89; P < .001; **Fig. 5B**). Within TBS+ regions, we observed that those regions exhibiting any form of plasticity had higher PLV (Z = 3.29; P < .001), CCEP amplitude (Z = -2.12; P = 0.03) and were closer to the stimulation site (Z = 4.22; P = <.001; **Fig. 5C**). In addition, in differentiating types of plasticity (within train, across train or both), PLV (Kruskal Wallis test; Chisqr = 6.8; P = 0.03), CCEP amplitude (Chisqr = 16.2; P < .001) and distance to stimulation site (Chisqr = 17.7; P < .001) were significantly different across these channel groups (**Fig. 5D**). Of note, sites showing across train plasticity were further away from the stimulation site than sites exhibiting within train plasticity (**Fig. 5D**). We obtained additional baseline metrics including CCEP duration and resting voltage correlations (see **Methods**). TBS+ channels had longer induced CCEPs after SPES at the same site (Z = -3.3; P < 0.001) and higher voltage correlations (Z = -6.0; P < 0.001; **Supplementary** Fig. 3). Channels with any type of plasticity had higher voltage correlations (Z = -3.3; P < .001) but no difference in CCEP response duration. Lastly, among the types of plasticity, CCEP duration was different across the types of response plasticity (Chisqr = 9.8; P = 0.007), but not voltage correlations (**Supplementary** Fig. 3).

**Figure 5:**
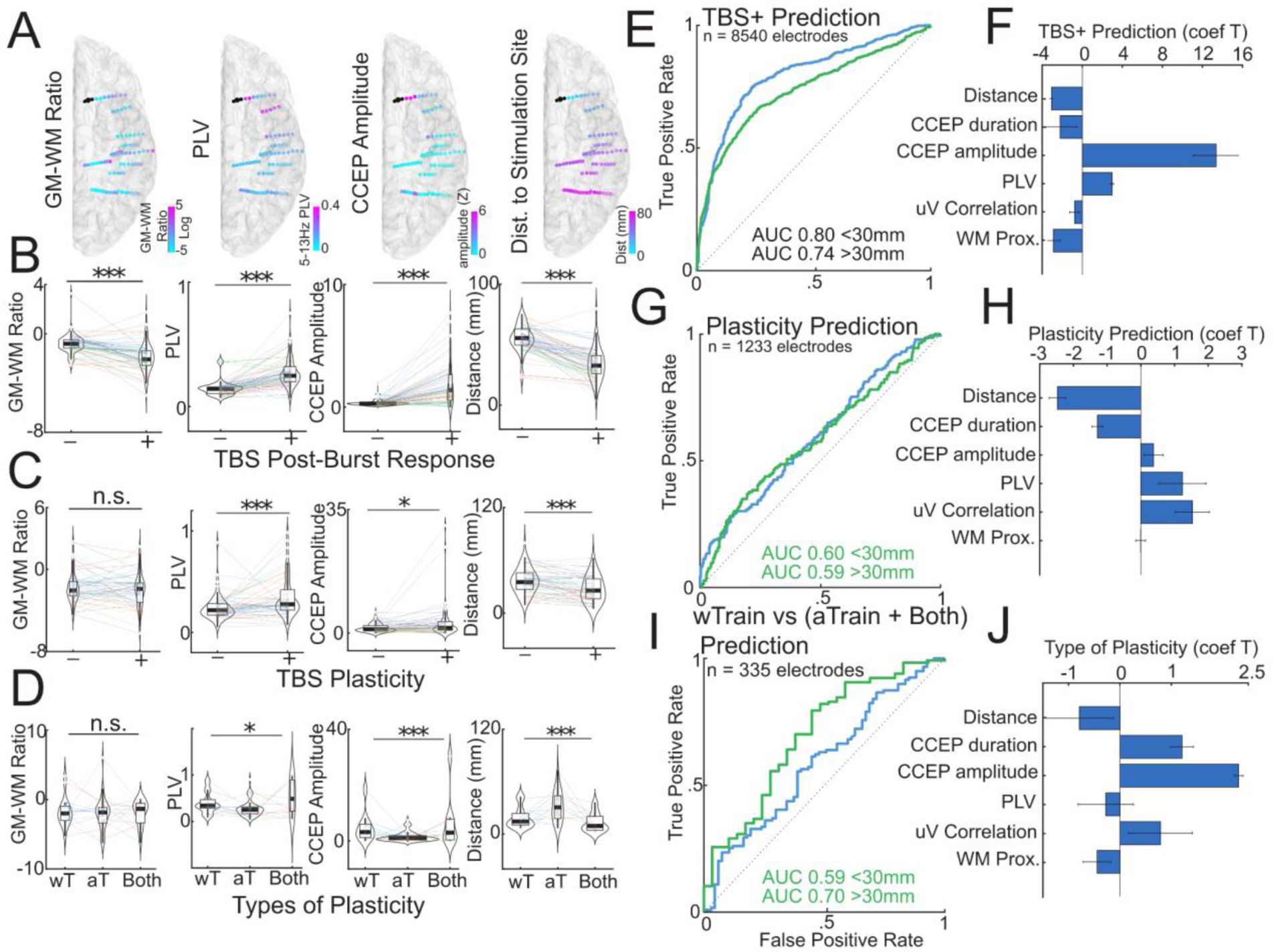
Structural, resting and effective connectivity at rest predict sites of TBS-evoked responses and plasticity. **(A)** Exemplar brains from a participant depicting variations in four baseline characteristics: 1) gray matter to white matter proximity ratio for a given channel in log-scale, 2) phase locking value (PLV) to the stimulation site for a given channel, 3) CCEP amplitude for a given channel and 4) Euclidean distance to the stimulation site for a given channel. **(B)** Differences in baseline characteristics in channels with and without significant TBS response. **(C)** Differences in baseline characteristics in channels with and without response plasticity. **(D)** Differences in baseline characteristics in channels with different types of response plasticity. Significant differences were observed for PLV, CCEP amplitude and distance to stimulation site, but not for gray matter to white matter proximity ratio. **I** Receiver Operating Characteristic (ROC) curves derived from baseline characteristics used to predict presence of a significant TBS post-burst response at a given channel. The curves are further stratified by channels either closer than or further from 30mm of the stimulation site. **(F)** The mean T-statistic for baseline features used in models to construct the ROC curves. Note CCEP amplitude is the most significant predictor for TBS+ channels. **(G)** ROC curves for prediction of plasticity using only TBS+ channels. **(H)** The mean T-statistic for baseline features used in models to construct the ROC curves. Note distance to stimulation site is the most significant predictor for channels with plasticity. **(I)** ROC curves for prediction of types of plasticity using only channels that demonstrated plasticity. **(J)** The mean T-statistic for baseline features used in models to construct the ROC curves. Note CCEP amplitude is the most significant predictor for type of plasticity.

We next asked if structural and functional measures can be used to predict the spatial distribution of TBS responses (**Fig. 5E-J**). We stratified the recording channels into local (<30mm) and distant (>30mm) groups relative to the stimulation location to take into account effects of volume conduction and as local vs distant sites may have differential responses ^69,85^. We constructed multivariate logistic regressions models with a ten-fold cross validation scheme using all computed structural and functional measures. We found that local TBS+ channels were predicted more reliably than distant TBS+ channels (**Fig. 5E**; Local AUC 0.80, 95%CI: 0.79-0.81; Distant AUC 0.74 95% CI: 0.72-0.76). CCEP amplitude had the largest feature coefficient T-statistic for prediction of TBS+ channels (**Fig. 5F**). Note the T-statistic is positive indicating that larger CCEP amplitude is associated with TBS+ prediction. We also generated additional predictive models using the individual features (**Supplementary** Fig. 4A). CCEP amplitude outperformed all other features in prediction of TBS+ channels (Local AUC 0.75, 95%CI: 0.73-0.86; Distant AUC 0.76 95% CI: 0.74-0.79). Limiting the selection to only TBS+ channels, we found that whether or not a channel undergoes plasticity can be predicted with above chance discrimination, whether local or distant (**Fig. 5G**; Local AUC 0.60 95%CI: 0.58-0.63; Distant AUC 0.59 95% CI: 0.56-0.62). Distance to stimulation site exhibited the highest feature coefficient T-statistic for predicting plasticity. Note the T-statistic is negative for distance indicating that decreasing distance to stimulation site is associated with plasticity. Distance to the stimulation site remains a leading feature in predicting sites with any plasticity compared to other individual features (**Supplementary** Fig. 4B; Local AUC 0.62, 95%CI: 0.59-0.66; Distant AUC 0.60 95% CI: 0.56-0.65). Lastly, in the subset of channels that exhibited plasticity, we found that these baseline measures can also be used to classify the *type* (within vs both types) of plasticity (**Fig. 5I**; Local AUC 0.59 95%CI: 0.52-0.66; Distant AUC 0.70 95%CI: 0.60-0.79). CCEP amplitude again had the largest feature coefficient T-statistic for predicting the type of plasticity (**Fig. 5J**). CCEP amplitude also outperforms other features on predicting plasticity when using individual features (**Supplementary** Fig. 4C; Local AUC 0.60, 95%CI: 0.50-0.71; Distant AUC 0.71 95% CI: 0.63-0.78). Therefore, measures such as distance to the stimulation site and connectivity can be used to predict whether we observe TBS responses as well as short-term plasticity at different time scales.

## DISCUSSION

We used direct electrical stimulation (DES) to identify neural responses to spaced, intermittent theta-burst stimulation (TBS) delivered across 29 sites in ten individuals. First, we characterized the effects of TBS by evaluating voltage deflections in response to each burst, focusing on temporal changes that may signify a form of short term plasticity ^58,59^. Second, we identified neuromodulatory effects of TBS resembling short-term plasticity such as facilitation and habituation that varied with dosage and stimulation. Finally, we examined the relationship between these effects to underlying anatomical and functional connectivity.

We initially confirmed the reliable elicitation of evoked responses from the theta-frequency bursts. These post-burst evoked responses were detectable in 14.4% of regions and were amplified with increased stimulation currents. Successive application of TBS bursts modulated these evoked responses in amplitude over time, indicating short-term plasticity occurring both within and across trains. Among the regions exhibiting post-burst responses, 27% demonstrated some form of short-term plasticity, with 19% exhibiting across-train changes, 5% within-train changes, and 2.5% experiencing both. Interestingly, across-train but not within-train plasticity was amplified with increasing stimulation current. Furthermore, leveraging baseline structural and resting functional connectivity profiles enabled accurate prediction of TBS response locations and their changes across bursts and trains, surpassing chance level. TBS responses exhibited high predictability (AUC = 0.75-0.80) whereas changes in TBS responses were less predictable (AUC = 0.60-0.70). Factors such as CCEP amplitude and anatomical distance between the recording and the stimulation sites played crucial roles in predicting both TBS responses and their alterations within and across trains.

Inherent to any iEEG study are certain limitations, including the clinical constraints of sparse sampling of brain tissue with sEEG and the restricted exploration of stimulation parameter due to time and safety constraints. While variability in stimulation location across patients poses a challenge, it also enriches our findings by offering generalizability. Our study focused on one type of patterned stimulation due to time constraints. Despite these limitations, our study provides valuable insights into the effects of TBS through direct electrical stimulation within the brain, leveraging high spatiotemporal resolution intracranial recordings. This is particularly important considering the field’s heavy reliance on noninvasive techniques like functional magnetic resonance imaging (fMRI) and electroencephalography (EEG), which offer limited temporal or spatial resolution. Yet, the temporal resolution of fMRI is poor (2-3 seconds), while scalp EEG has good temporal resolution (e.g., 10ms) but suffers from poor spatial resolution, insufficient to identify the regional effects of patterned stimulation. By capturing brain responses with fine temporal precision at precise intracranial localization, we aimed to elucidate how TBS-patterned direct electrical stimulation influences neural activity in the human brain.

Understanding how TBS-patterned direct electrical stimulation induces plasticity in humans, remains critical but limited in humans in vivo, particularly in the frontal lobe. Previous studies in epilepsy patients evaluated the effects of TBS in the sensorimotor cortex, revealing modifications in beta-frequency coherence ^56^. While other iEEG studies have examined TBS-induced oscillatory changes, notably increasing theta band power ^12,100^, the voltage effects of TBS and the dynamics of these neural responses within and across stimulation trains remain unexplored. This is critical as these features – voltage changes after stimulation – underlie neuroplasticity studies in animal models ^57–60^. Our results indicate the reliable quantification and temporal tracking of evoked response immediately following each high-frequency burst (*TBS* response). The neuronal origin of this response is likely complex, reflecting an interaction of both monosynaptic and polysynaptic responses^50,101^. The observation that these post-burst responses dynamically change over time indicate a form of short-term plasticity, potentially linked to underlying neuronal responsiveness or synaptic modifications, though further investigations would be needed to answer this question.. Our prior work with 10Hz direct brain stimulation demonstrated that voltage changes observed during the stimulation protocol itself is highly predictive of subsequent post-stimulation short-term changes (on the scale of minutes) in network connectivity^43,95^. In this study, we tested how varying different stimulation parameters such as current amplitude and stimulation site impact both the immediate post-burst response as well as changes in the post-burst response (short-term plasticity). We found that stronger TBS (higher amplitude) resulted in broader engagement of cortical regions and drove changes in post-burst responses over time. In addition, each stimulation site exhibited a distinct spatial profile of TBS-responsive brain regions, emphasizing the importance of understanding of how stimulation parameters modulate TBS responses to each burst as well as changes in the TBS response across bursts and trains (plasticity).

The type of short-term plasticity observed occurring between bursts (within train) and across trains could in the future be used as an acute indicator to optimize TBS treatment. Specifically, this indicator could be optimized in closed-loop fashion to maximize plasticity effects after a single treatment, metaplasticity effects after multiple treatments, and eventual clinical effects ^102–104^. Of course, the fact that the within train and across train plasticity is occurring at two different time scales (∼0.100 sec vs 20 seconds) could indicate different mechanisms of induced plasticity such as differentiating facilitation and habituation (as with within-burst plasticity) versus response changes shown here which may be on the order of tens of seconds ^57,59,105^. Granted, only a limited range of stimulation parameters were explored in this study, as we prioritized sampling across stimulation sites and current amplitudes. Future studies will evaluate other parameters of TBS including TBS patterns used to treat depression (3 pulses / burst, 10 bursts / train, 20 trains / dose), and other less explored parameters including burst pulse number and frequency, inter-burst interval, number of bursts / train, inter-train interval, and number of trains. Notably, future studies will be needed to test if these findings translate to the FDA-cleared TBS protocol for depression (triplet at 50Hz, 10 bursts / train @ 5Hz, 8s inter-train interval). For instance, one prediction is that the shorter temporal spacing (8 sec vs 20 sec) or more trials could induce larger short term plastic changes, for instance, though it would be interesting to determine whether some brain regions are more evidently plastic than others. Relevant to this point, TMS sessions of intermittent TBS are usually repeated every 24 hours though recent evidence indicates applying multiple treatments per day can achieve a faster therapeutic effect, in as short as just a few days ^19^. Further, the neural effects of other non-TBS forms of patterned stimulation (e.g. beta burst and other non-burst patterned stimulation) will be evaluated, particularly if these other types of stimulation could uncover other forms of plasticity in the brain.

Our observation that repeated TBS trains led to dynamic changes in voltage responses could help explain why repeated sessions of TMS-delivered TBS leads to improved clinical outcome in depression and other psychiatric disorders ^106–108^. This type of state-dependent accumulated brain changes that may result from spaced and repeated stimulation is referred to as metaplasticity and is well-known in non-human models ^102–104^. Indeed, intermittent TBS – delivering repeated and spaced TBS trains (as in this study) –has been shown to produce neural changes that could reflect plasticity while continuous TBS (no spacing between trains) did not – an effect which could be NMDA-receptor dependent ^14,68,109^. In other words, intermittent TBS could be more conducive to triggering plastic changes in the human brain.

A major finding in this work is that functional, effective, and structural connectivity all contributed to an ability to predict the post-burst response as well as the type of plasticity induced by TBS. This is consistent with multiple past studies which have indicated that underlying connectivity shapes stimulation responses, though few of those studies examine whether this connectivity can predict changes in the brain ^13,43,50,69–75^. Since TBS responses and even plasticity could be predicted based solely on pre-TBS metrics, this opens up several exciting potential applications. In the future one could select the optimal stimulation target by considering the pre-treatment structural and functional connectivity of a set of possible stimulation sites. The two most predictive features we observed were 1) the distance to the stimulation site and 2) CCEP amplitude. Translating this to the non-invasive space, diffusion-tensor imaging (DTI) could be employed to define anatomical tracts, while single pulse stimulation using TMS and recording with EEG can be used to index effective connectivity ^110–115^. Based on these two parameters, a gradient of how likely the target brain area would undergo plasticity from stimulation at different locations can be constructed, prior to any stimulation. Further, these TBS results shown here may be useful in predicting what circuits could be engaged with stimulation in certain regions non-invasively, such as the result that DLPFC stimulation largely induced ACC responses with TBS.

In sum, we observed that direct electrical intermittent TBS induces reliable immediate voltage responses which are modified by the history of stimulation in a predicable subset of brain regions based on functional connectivity. These results offer further insight in how patterned neurostimulation alters neural activity, and how stimulation therapies can be optimized to maximize neuroplasticity and, ultimately, therapeutic effects.

## Acknowledgments

We would like to especially thank the patients for participating in the study. We would like to thank Dan Soper, Jessica Chang, and Pariya Salami for help in data collection.

