## SupplementalData for "Theta-burst direct electrical stimulation remodels human brain networks"

**Supplemental Table 1. Participant designations and information**

| Participants | Diagnosis | Implant hemi-sphere | Hand-edness | Previous surgery | TBS current (mA) | No. of TBS sites |
|---|---|---|---|---|---|---|
| sub-1a6g* | epilepsy, hippocampal sclerosis, but underlying etiology likely rupture of an arteriovenous malformation | unilateral right | Right | resection | 1 & 2 | 4 |
| sub-2j8e | left mesial temporal lobe epilepsy of unknown etiology | unilateral left | Right | none | 1 & 2 | 2 |
| sub-4u3s | cortical epilepsy of unknown etiology | unilateral left | Right | none | 1 & 2 | 2 |
| sub-5o1r | frontal lobe epilepsy with unknown etiology | bilateral | Right | none | 1 & 2 | 4 |
| sub-7k6q | mild granule cell dispersion and focal pyramidal cell loss in hippocampus, hippocampal epilepsy etiology unknown | bilateral | Right | none | 1 & 2 | 2 |
| sub-7q1t* | bitemporal epilepsy, non-lesional focal onset seizures | bilateral | Right | RNS | 1 & 2 | 4 |
| sub-7u9e* | epilepsy related to right hemispheric stroke | unilateral right | Right | none | 1 & 2 | 4 |
| sub-8d4f* | left mesial temporal lobe epilepsy with mesial temporal lobe sclerosis | unilateral left | Right | none | 1 | 4 |
| sub-9d4z | epilepsy secondary to astrocytoma | unilateral left | Left | resection | 1 & 2 | 2 |
| sub-9u0z | bi-hippocampal epilepsy, etiology unknown | bilateral | Right | none | 1 & 2 | 2 |

***** - participant data where single pulse electrical stimulation has been reported in previous publications and is shared on data repositories [1,2]

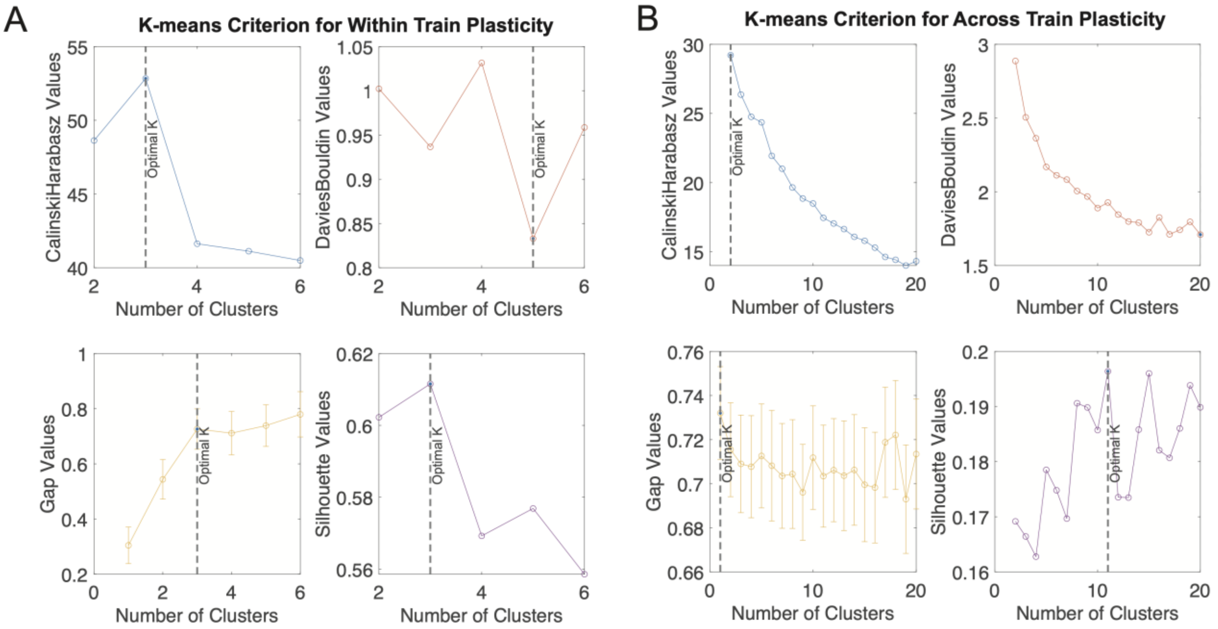


**Supplementary Figure 1: K-means cluster criterion for optimal number of clusters in post-burst patterns over time. (A)** Criterion evaluation for pattern of within train response modulation. Calinski-Harabasz, Gap and Silhouette values converged at K = 3 as an optimal number of clusters. DaviesBouldin values indicated optimal K = 5. **(B)** Criterion evaluation for pattern of across train response modulation. No consensus was achieved amongst Calinskin-Harabasz, DaviesBouldin, Gap and Silhouette values for optimal number of clusters.

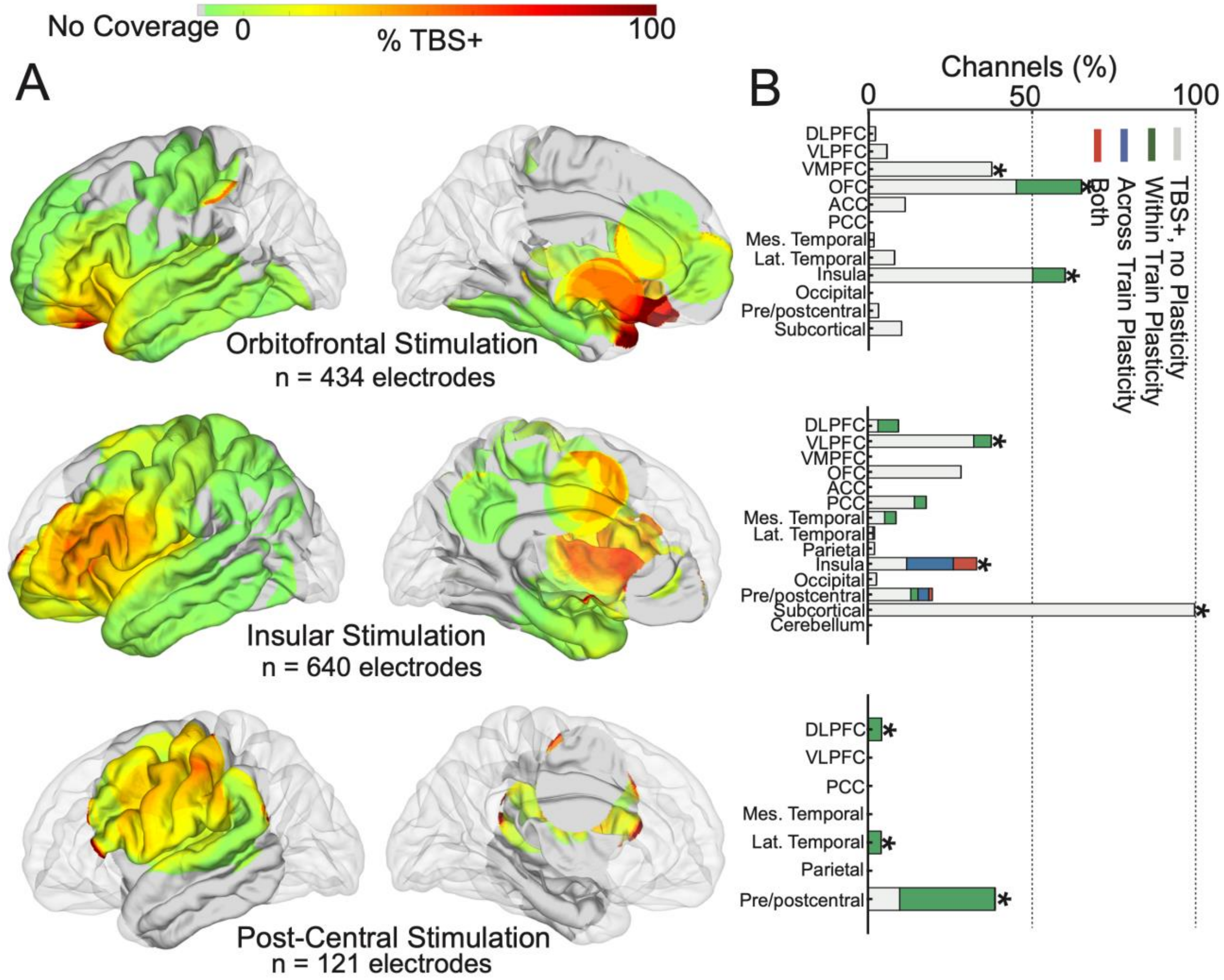


**Supplementary Figure 2: Spatial specificity of theta-burst stimulation responses and plasticity for sparsely stimulated sites. (A)** Surface heatmap and **(B)** bar chart depicting the percentage of TBS+ local electrodes for a particular stimulation site (gray), within train response modulation (blue), across train response modulation (green) and both types of response modulation (red). Top three sites for each stimulation location are noted with asterisk. DLPFC: dorsolateral prefrontal cortex; VLPFC: ventrolateral prefrontal cortex; VMPFC: ventromedial prefrontal cortex; OFC: orbitofrontal cortex; ACC: anterior cingulate cortex; PCC: posterior cingulate cortex.

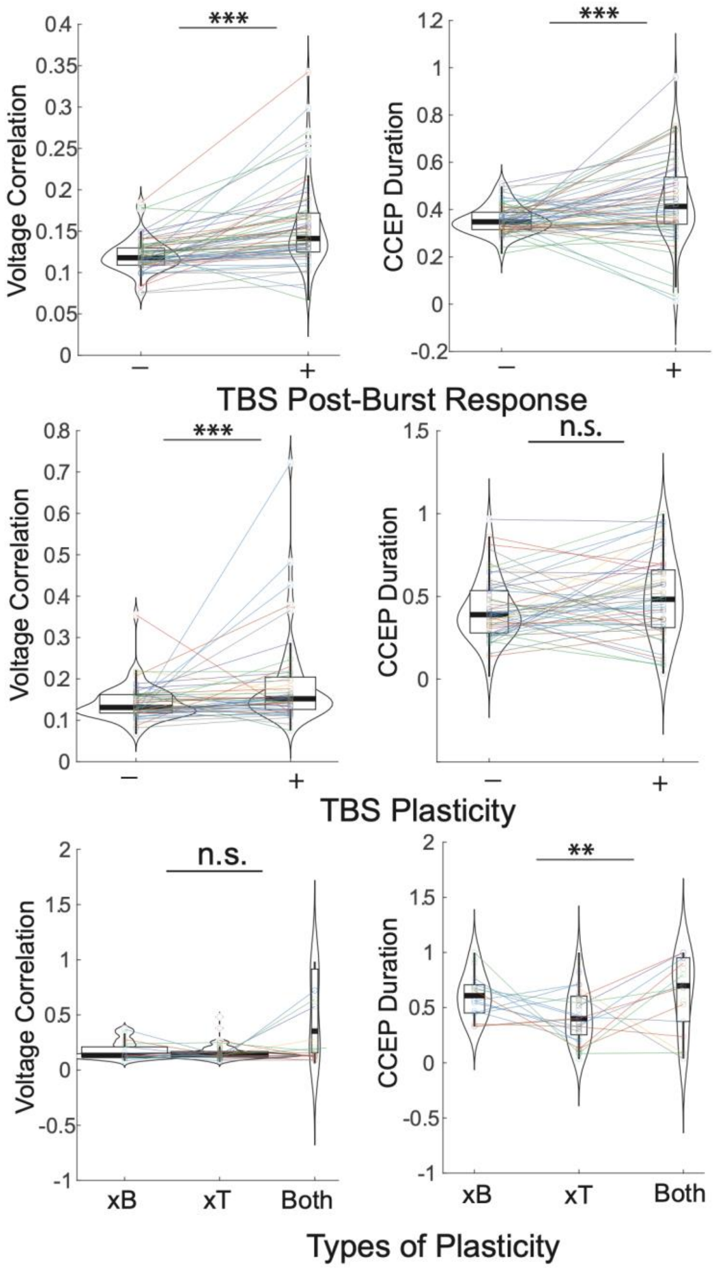


**Supplementary Figure 3: Additional resting characteristics and relationship to TBS responses.** Voltage correlation at rest and CCEP duration were compared across the following channel groups: TBS+ vs TBS-, plasticity vs no plasticity, and different types of plasticity (within train, across train, or both).

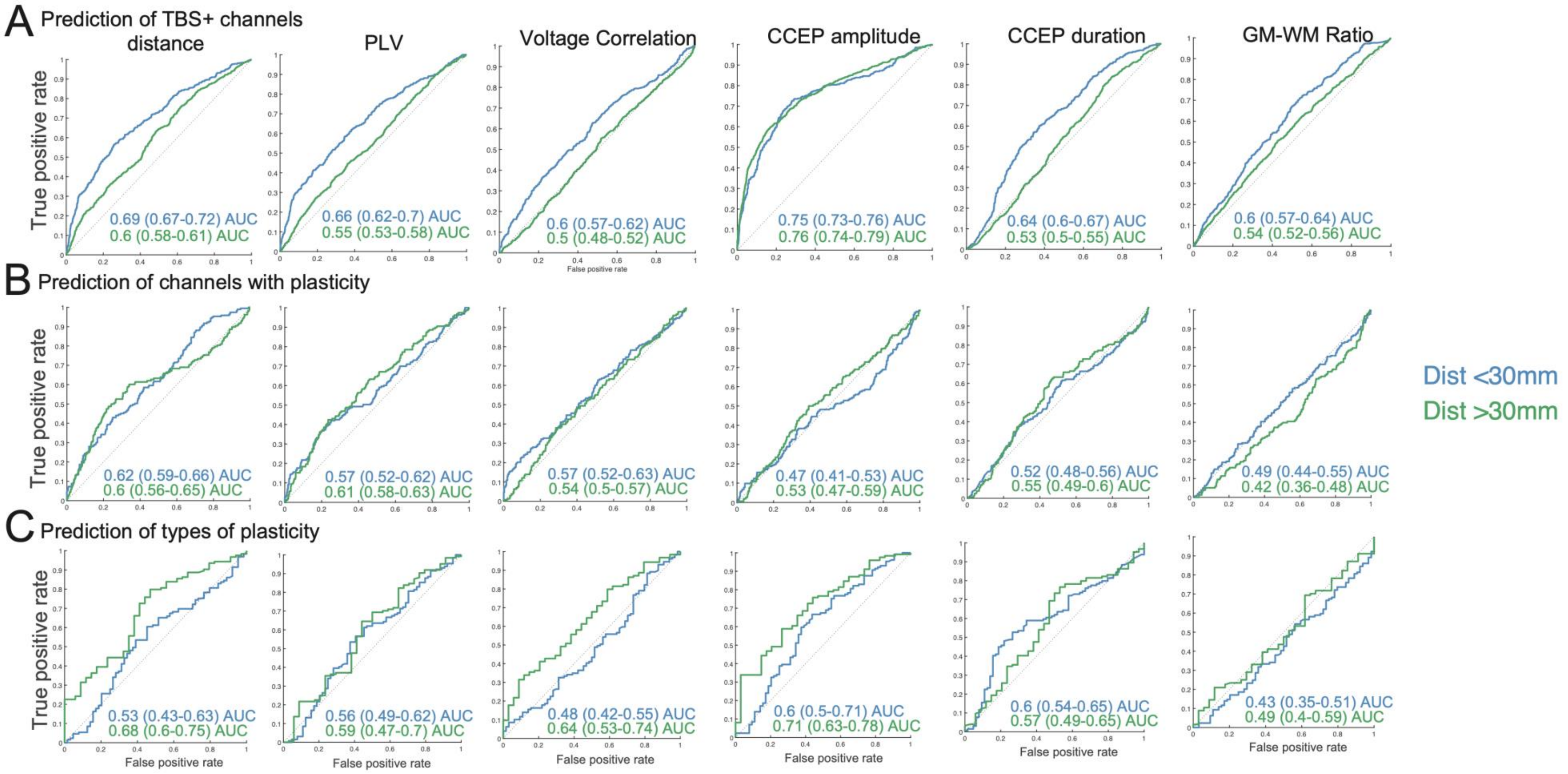
A Prediction of TBS+ channels
distance
PLV
Voltage Correlation
CCEP amplitude
CCEP duration
GM-WM Ratio
True positive rate
False positive rate
0.69 (0.67-0.72) AUC
0.6 (0.58-0.61) AUC
0.66 (0.62-0.7) AUC
0.55 (0.53-0.58) AUC
0.6 (0.57-0.62) AUC
0.5 (0.48-0.52) AUC
0.75 (0.73-0.76) AUC
0.76 (0.74-0.79) AUC
0.64 (0.6-0.67) AUC
0.53 (0.5-0.55) AUC
0.6 (0.57-0.64) AUC
0.54 (0.52-0.56) AUC
B Prediction of channels with plasticity
0.62 (0.59-0.66) AUC
0.6 (0.56-0.65) AUC
0.57 (0.52-0.62) AUC
0.61 (0.58-0.63) AUC
0.57 (0.52-0.63) AUC
0.54 (0.5-0.57) AUC
0.47 (0.41-0.53) AUC
0.53 (0.47-0.59) AUC
0.52 (0.48-0.56) AUC
0.55 (0.49-0.6) AUC
0.49 (0.44-0.55) AUC
0.42 (0.36-0.48) AUC
C Prediction of types of plasticity
0.53 (0.43-0.63) AUC
0.68 (0.6-0.75) AUC
0.56 (0.49-0.62) AUC
0.59 (0.47-0.7) AUC
0.48 (0.42-0.55) AUC
0.64 (0.53-0.74) AUC
0.6 (0.5-0.71) AUC
0.71 (0.63-0.78) AUC
0.6 (0.54-0.65) AUC
0.57 (0.49-0.65) AUC
0.43 (0.35-0.51) AUC
0.49 (0.4-0.59) AUC
Dist <30mm
Dist >30mm

**Supplementary Figure 4: Individual resting feature prediction of TBS responses. (A)** Prediction of TBS+ channels using individual features, including distance to stimulation site, phase locking value, resting voltage correlation, CCEP amplitude, CCEP duration and gray matter to white matter ratio. Each prediction was further stratified into prediction within either local (<30mm from stimulation site) or distal (>30mm from stimulation site) channels. **(B)** Prediction of channels with response modulation using individual features as in (A). **(C)** Prediction of different types of response modulation using individual features as in (A).

1. Paulk, A. C. *et al.* Local and distant cortical responses to single pulse intracranial stimulation in the human brain are differentially modulated by specific stimulation parameters. *Brain Stimulat.* **15**, 491–508 (2022).
2. Zelmann, R. *et al.* Differential Cortical Network Engagement During States of Un / Consciousness in Humans. *Res. Sq.* 1–28 (2022) doi:https://doi.org/10.21203/rs.3.rs-2006868/v2.
